# Role of ERK activation in *H. pylori*-induced disruption of cell-cell tight junctions

**DOI:** 10.1101/2020.09.03.276600

**Authors:** Amita Sekar, Bow Ho

## Abstract

**Background:** Tight junctions, a network of claudins and other proteins, play an important role in maintaining barrier function and para-cellular permeability. *H. pylori*, the major etiological agent of various gastroduodenal diseases, is known to cause tight junction disruption. However, the molecular events that triggered cell-cell tight junction disruption in *H. pylori*-infected cells, remain largely elusive.

**Materials and Methods:** Trans-epithelial electrical resistance (TEER) and FITC-Dextran permeability measurement were performed to determine the barrier function in *H. pylori* 88-3887-infected polarized MKN28 cells. For visualization of tight junction protein localization, immunofluorescence and immunoblotting techniques were used. To examine the role of ERK activation in tight junction disruption, U0126, a MEK inhibitor, was employed. To further support the study, computational analyses of *H. pylori*-infected primary gastric cells were carried out to decipher the transcriptomic changes.

**Results:** The epithelial barrier of polarized MKN28 cells when infected with *H. pylori* displayed disruption of cell-cell junctions as shown by TEER & FITC-dextran permeability tests. Claudin-4 was shown to delocalize from host cytoplasm to nucleus in *H. pylori*-infected cells. In contrast, delocalization of claudin-4 was minimized when ERK activation was inhibited. Interestingly, transcriptomic analyses revealed the upregulation of genes associated with cell-junction assembly and ERK pathway forming a dense interacting network of proteins.

**Conclusion:** Taken together, evidence from this study indicates that *H. pylori* regulates ERK pathway triggering cell-cell junction disruption, contributing to host pathogenesis. It indicates the vital role of ERK in regulating key events associated with the development of *H. pylori*-induced gastroduodenal diseases.

## Introduction

Gastric mucosal barrier, which is composed of thick mucus layer and polarized epithelial cells, confers protection to the inner layers of the stomach from the hostile acidic environment^[1][2]^. Despite being a single layer of epithelial cells, it provides a sturdy wall against the highly corrosive contents of the gut, maintaining homeostasis. Polarized gastric epithelial cells harbour distinct domains of which the lateral domains comprise cell-cell junctions and account for establishing cell-cell contact and cell adhesion to the extracellular matrix^[3]^. Cell-cell tight junctions found at the apical side are responsible for the tight packaging of the cells and maintaining paracellular permeability. Tight junctions are composed of transmembrane proteins such as Junction Adhesion Molecule (JAM), Occludins, Claudins and cytoplasmic connector protein Zona Occludens (ZO-1). Together, these proteins are closely associated to form a tight junction assembly^[4]^ that maintain the cell barrier integrity.

Focussed research on claudins from the time of its discovery in 1998 has garnered mounting evidence that these junction proteins form the primary physical basis of tight junction assembly^[5]^. The homophilic and heterophilic interactions among different claudins as well as its association with the other components of tight junction complex is pivotal in ensuring proper functioning of the cell-cell tight junctions^[6]^. Till date, 27 claudins have been reported and they have been found in tissue specific combinations thereby conferring tissue-specific barrier properties^[9]^. Phosphorylation of claudins has been reported to play a vital role in paracellular permeability. Various kinases like myosin light chain kinase (MLCK) and protein kinase A (PKA) have been shown to be involved in the phosphorylation of claudins^[10]^.

Kwon (2013) observed aberrant expression of cell-junction associated proteins in clinically isolated tumour samples^[11]^. Similarly, moderate to high staining of claudin-4 was correlated to decreased survival in gastric adenocarcinoma tissues suggesting a strong link between expression of tight junction proteins and cancer^[12]^. The altered expression and localization of another tight junction protein, ZO-1, was observed in metastatic pancreatic ductal adenocarcinoma tissue^[13]^. In contrast, reduced expression of several claudins has also been closely linked to tumorigenesis. Claudin-1 and -7 have been found to be deregulated in breast carcinoma while downregulation of claudin-4 was observed in hepatocellular carcinoma^[9]^. These studies highlight the importance of expression and localization of cell-junction protein in maintaining host cell integrity. Substantial evidence has thus led us to consider cell junction proteins as potential biomarkers in various disease conditions, mostly in cancer.

Tight junction disruption has been reported to be caused by various microbes. In the course of pathogen infection, inflammation and other host responses have also shown to cause internalization of claudins which has a direct effect on the host cell homeostasis. Bacteria such as *Listeria monocytogenes, Campylobacter jejuni* and *Escherichia coli* have been shown to invade the host causing tight junction disruptions^[14][15][16]^. However, the mechanism by which these pathogens affect the cell-cell junction proteins to cause a physiological imbalance in host cells remains largely elusive.

*Helicobacter pylori*, an extensively studied gram negative gut pathogen has been highly associated with various gastro-duodenal diseases including gastric cancer^[17][18]^. More than half of the world’s population are infected with this gut bacterium^[19]^. The bacterium harbors an array of virulence factors which aid in colonization of the gut. Bacterial proteins such as cytotoxin associated gene A (CagA) and vacuolating cytotoxin A (VacA) have been determined to be of high importance in *H. pylori* pathogenesis^[20]^. *H. pylori* has been reported to translocate CagA via the type IV secretion system (T4SS) into the host cells triggering events such as IL-8 induction, cytoskeletal rearrangement and many other host responses^[21][22]^. Intriguingly, the CagA-T4SS system was found to interact with basolateral integrins to translocate the virulence factor into the host^[23][24]^. These findings have led to speculate that *H. pylori* could induce cell-cell junction disruption prior to accessing the basolateral integrins in order to translocate CagA into the host^[25]^. In 2003, Ameiva *et al* observed that CagA could play a role in recruiting ZO-1 to the sites of bacterial attachment and cause cytoskeletal rearrangement^[26]^. Factors such as high temperature requirement A (HtrA) have been reported to aid in *H. pylori*–induced cell junction disruption^[27]^. In other studies, *H. pylori* urease has been implicated in causing tight junction disruption by targeting claudin-4 and -5^[28][29]^. Even though strong evidence suggests the involvement of *H. pylori* in tight junction disruption, the exact mechanism by which the bacterium induces tight junction disruption remains unclear.

ERK is a major transcription factor involved in the expression of many crucial proteins determining the fate of host cell. *H. pylori* has been shown to activate ERK triggering many host response pathways^[30][31][32][33][34]^. Several reports have strongly implicated the role of activated ERK in regulating cell junction proteins, particularly claudins, and in disrupting cell-cell barrier function. Wang *et al*., (2004) demonstrated that activation of ERK1/2-MAPK pathway was found to disrupt tight junctions in human corneal epithelial cells^[35]^. Interestingly, Aggarwal *et al* in 2011 reported that knockdown of ERK1/2 enhanced tight junction integrity while U0126 (ERK kinase inhibitor) attenuated the disruption initiated by the activation of EGFR^[36]^. In contrast, Ray *et al* in 2007 revealed that MEK1/ERK is highly associated with maintaining cell-cell contacts^[37]^. A recent published study observed that inhibition of ERK signaling pathway decreased the expression of claudin-2 in lung adenocarcinoma cells^[38]^ and similar results were also observed in Madin-Darby Canine Kidney cells (MDCK) I and II cells^[39]^. Although several studies have explored the role of *H. pylori*-induced ERK activation, there is a knowledge gap regarding its effect on tight junction proteins and barrier function disruption.

Our study aims to deduce the role of ERK activation in *H. pylori*-induced barrier function disruption. We present data showing the role of *H. pylori*-induced ERK activation in delocalizing claudin-4 from the tight junctions leading to the disruption of barrier function in MKN28 cells. This study also provides supporting data from transcriptomic analysis of *H. pylori*-infected primary cells showing significant regulation of tight junction and ERK signalling related gene expression.

## Methods

### Bacterial and in vitro cell cultures

*H. pylori* 88-3887 strain was used in this study. Bacteria were grown in 5% chocolate blood agar plates at 37°C in a 10% CO_2_ incubator (Forma Scientific, USA) for two days before being harvested and suspended in phosphate buffered saline (PBS). A standard curve was generated based on the correlationship between the optical density readings of bacterial suspensions and bacterial count using plate count.

Polarized MKN28 cells were used as the study model as these cells have been shown to form intact cell-cell junctions^[29]^. The cells were cultured and maintained in RPMI 1640 with 2.05 mM L-glutamine (HyClone, USA) supplemented with 10% Fetal Bovine Serum (HyClone, USA) and incubated at 37°C in a 5% CO_2_ incubator (Forma Scientific, USA). The polarized cells were grown to above 90% confluency to ensure formation of intact cell junctions.

### Materials

FITC-Dextran-4kDa used in permeability assay was purchased from Sigma-Aldrich, USA. Mouse monoclonal anti-Claudin-4 (1:100, Novex, Life Technologies, USA) and rabbit polyclonal anti-claudin-4 (1:1000, Santa Cruz Biotechnology, USA) were used in immunofluorescence and immunoblotting experiments respectively. Rabbit polyclonal anti-*H. pylori* (1:250) was from Dako, Denmark and mouse monoclonal anti-*H. pylori* (1:250) was from Thermo Scientific, USA. Mouse monoclonal Beta-actin (1:3000) from Cell Signalling Technologies, USA was used as marker for whole cell protein immunoblotting experiments. Rabbit polyclonal p-cadherin (1:1000, Santa Cruz Biotechnology, USA), Rabbit polyclonal beta-tubulin (1:1000, Cell Signalling Technologies, USA) and Rabbit polyclonal anti-Lamin A/C (1:1000, Cell Signalling Technologies, USA) were used as markers for membrane, cytoplasmic and nuclear protein respectively. ERK activation was assessed using rabbit polyclonal anti-p-ERK (1:500, Cell Signalling Technologies, USA). Polyclonal goat anti-mouse IgG/HRP and Polyclonal goat anti-rabbit IgG/HRP (1:2000) were from Dako, Denmark. Fluorescent conjugated secondary antibodies Cy3 goat anti-mouse IgG (H+L) (1:100), Cy3 goat anti-rabbit IgG (H+L) (1:100), Alexafluor 488 F(ab’)2 fragment of goat anti-rabbit IgG (H+L) (1:100) and Alexafluor 488 goat anti-mouse IgG (H+L) (1:100) were all purchased from ThermoFisher Scientific, USA. U0126 (MEK inhibitor) was purchased from Cell Signaling Technologies, USA. For the inhibitor study, the cells were treated with 10uM U0126 (Tocris Biosciences, UK).

### Infection study

The confluent cells were infected with *H. pylori* 88-3887 at a multiplicity of infection (MOI) of 100:1. The cells were then incubated at 37°C in a 5% CO_2_ incubator for 24 hrs or various time points as required for the experiment. Post infection, the cells were washed 3 times with 1x PBS prior to any further downstream processing.

### Immunofluorescence studies

Cells were cultured on sterilized coverslips placed in 6-well plates (Greiner Bio-One, Austria) containing culture medium RPMI 1640 for 3-4 days. The confluent cells were treated with inhibitor and/or infected with *H. pylori* while uninfected cells served as control. At the required time points, the cover slips containing the cells were withdrawn and washed twice with 1 x PBS before being fixed using 3.7% formaldehyde. Following this, the cells were permeabilized using 3.7% formaldehyde and 0.02% Triton X-100 (Sigma-Aldrich, USA). Blocking was carried out with 2% Bovine Serum Albumin (BSA) (Sigma-Aldrich, USA) for 2 hours, prior to incubating with primary antibodies at 4°C overnight. The cells were then washed thrice with 1 x PBS for 5 mins each prior to incubating with fluorescent conjugated secondary antibodies for 2 hours at room temperature. The coverslips containing the immuno-stained cells were washed thrice with 1 x PBS then mounted onto glass slides using mounting media with DAPI (Vector Laboratories, Inc., USA). The stained cells were viewed using confocal laser scanning microscopy (CLSM) (Olympus FV3000, Japan).

3D image reconstruction: z-stack images were taken using CLSM and these images were loaded onto Imaris software (version 7.6, Bitplane) for reconstructing the images in a 3D plane. These images were used to analyse the delocalization of claudin-4 from the apical membrane.

### Transepithelial electrical resistance (TEER) measurement

Cells were cultured in 12-well plate hanging inserts (Greiner Bio-One, Austria) for 3-4 days to reach > 90% confluency. TEER readings were measured using the electrode supplied along with the MilliCell-ERS volt-ohm meter (Merck, USA). A blank insert with media alone was used as a negative control. MKN28 cells with TEER reading > 350 Ω.cm^2^ was considered optimal for the barrier function studies. TEER readings were measured for cells subjected to inhibitor treatment and/or infected with *H. pylori*. Uninfected cells served as control. The % baseline resistance was calculated using the following equation and the values were plotted as a graph. All experiments were performed in triplicates.

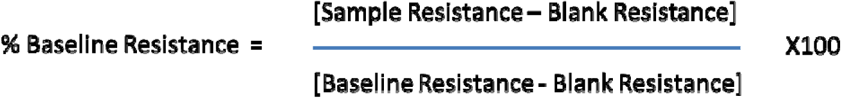

### FITC-Dextran permeability assay

MKN28 cells were cultured for 3-4 days to confluency in 12-well inserts and subjected to inhibitor treatment and/or infection with *H. pylori*. Uninfected cells served as control. The cells were washed twice with 1 x PBS before adding FITC-Dextran (1mg/ml) to the upper chamber. The FITC-Dextran that diffused into the basal chamber media was determined by measuring the fluorescence intensity at optical density 492nm (Tecan Infinite, Switzerland). All experiments were performed in triplicates.

### Protein extraction and western blot studies

Post infection cells were lysed using Triton-X 100 containing lysis buffer to extract whole cell proteins. Membrane and cytoplasmic protein fractions of cells infected with *H. pylori* for 24 hours were extracted using ProteoJet™ Membrane Protein extraction kit (Fermentas, Thermo Scientific, USA) according to the manufacturer’s protocol. Cytoplasmic and nuclear fractions were separated using NE-PER™ Nuclear and Cytoplasmic Extraction Reagents (Thermo Scientific, USA). The protein concentration of samples were estimated using Bradford Assay (Bio Rad, USA) and an equivalent of 15ug protein was loaded onto 12% SDS-Poly Acrylamide gel and subjected to protein electrophoresis. The samples were then electro-blotted onto PVDF membrane by wet transfer method. The membranes were blocked with 2% BSA prior to incubating with primary antibodies at 4°C overnight. The membranes were then washed thoroughly with 1xPBS with 0.1% Tween-20 (Sigma-Aldrich, USA) for 5 times of 5 mins each. The membranes were then incubated with HRP-conjugated secondary antibodies for 2 hours at room temperature. After thorough washing (5 x 5 mins), the membranes were developed using Pierce ECL western blotting substrate (Thermo Scientific, USA) and the bands were documented using ChemiDoc™ MP System (Bio-Rad Laboratories, USA).

### q-PCR studies

RNA was extracted from infected and uninfected cells using RNeasy Mini kit (Qiagen, USA) and quantitated spectrophotometrically. Specific primers for claudin-4 [Forward primer 5’-TGGGAGGGCTATGGATGAA-3’, Reverse Primer 5’-GCTTTCATCCTCCAGGCAGT-3’] and endogenous control GAPDH [Forward primer 5’-ATCTCCCCTCCTCACAGTTG-3’, Reverse Primer 5’-TGGTTGAGCACAGGGTACTT-3’] were synthesized by Sigma-Aldrich, USA. These primers were used to amplify the mRNA from the sample using Quantifast SyBR Green one –step RT-PCR kit (Qiagen, Netherlands) and the reactions were run using ABI 7500 real-time PCR instrument (Thermo Scientific, USA). The experiment was performed in triplicates and the relative quantitation values were analysed using 7500 software v2.0.6 (Thermo Scientific, USA) and represented as a graph.

### Bioinformatic analysis

Published RNA-Seq raw data of *H. pylori*-infected gastric primary cells were used for downstream analysis (Accession number: GSE55699)^[40]^. The published libraries were downloaded from Gene Expression Omnibus^[41]^.

The fastq format raw reads were subjected to quality control analysis using FASTQC (http://www.bioinformatics.babraham.ac.uk/projects/fastqc/). The sequencing files were considered to be of high quality when the following parameters were observed: (Mean Quality score-30 and above, Adapter contamination level-Below 0.01%, Duplication level-Below 80%, %A ∼%T, %G∼%C).

Mapping of the RNA-Seq libraries was performed using the splicing-aware STAR 2.4. The software ensured trimming of unmappable and low quality sequences. Reads with more than 2 mismatches and reads that map to more than one locus in the genome were filtered out. The mapped files were imported into SeqMonk 0.31.0 (http://www.bioinformatics.babraham.ac.uk/projects/seqmonk/) and the replicates and sample groups were assigned appropriately. Mapping QC was performed using “RNA-Seq QC Plot” option. Differential gene expression analysis was carried out using the in-built EdgeR statistics module^[42]^. Mapped files were then normalized using the log10 (RPKM) transformation option in RNA-Seq quantitation pipeline module. Heatmaps of the significant genes were generated using the “Per-probe normalized Hierarchical Clustering” option. Gene Ontology terms file (.bgo) and the human gene ontology annotation file were downloaded from the gene ontology consortium website (http://geneontology.org/). These files were imported to BiNGO^[43]^ application which is installed in Cytoscape 3.2.0^[44]^. List of significant genes was uploaded to BiNGO to identify the enriched biological processes. Gene lists of relevant biological processes were uploaded to STRING (http://string-db.org/) protein-protein interaction database^[45]^. The text mining option was disabled. The interaction network was downloaded in text format. The text file was uploaded to Cytoscape 3.2.0 for visualization.

### Statistical analysis

Student t-test (2-tailed, paired end) was used to calculate the p-value and determine statistical significance.

## Results

### H. pylori affects barrier function and impairs the localization of claudin-4 in MKN28 cells

Cell-cell junctions confer epithelial barrier function and play a critical role in permeability. Epithelial barrier function of polarized MKN28 cells were determined using Trans-epithelial electrical resistance (TEER) measurement and FITC-Dextran permeability assay. TEER readings taken 24 hours post-*H. pylori* infection were found to be significantly reduced (p-value = 0.009050856) in *H. pylori*-infected cells when compared to the uninfected control (Fig. 1A, left panel). Similarly, permeability of 4kDa FITC-Dextran was found to be higher in *H. pylori*-infected cells compared to uninfected cells (Fig. 1A, right panel). The study demonstrates *H. pylori*-induced barrier function disruption in *H. pylori*-infected MKN28 cells using these two techniques.

**Fig.1.**
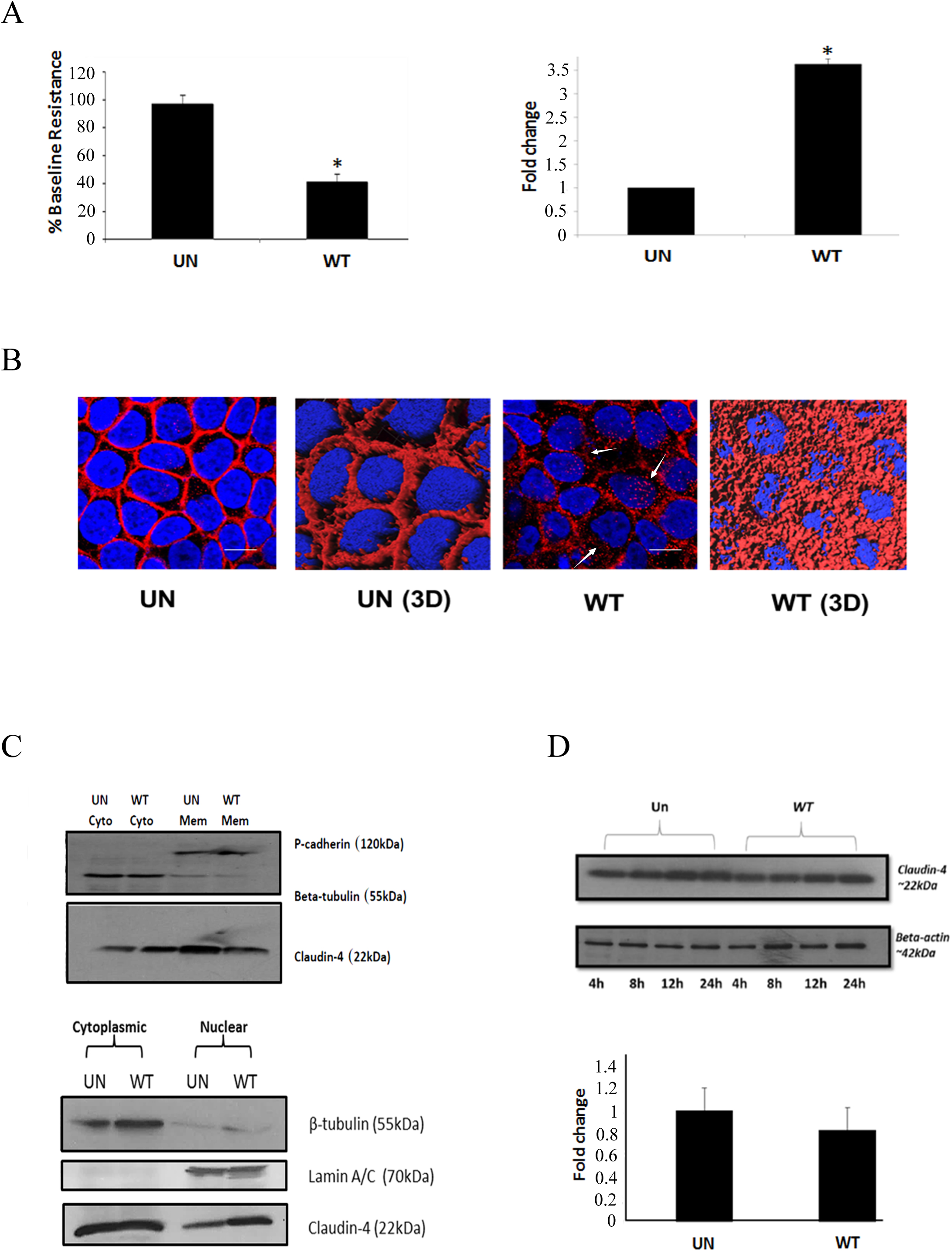
Cell-cell tight junction disruption and delocalization of claudin-4. MKN28 cells were infected with *H. pylori* 88-3887 for 24 hours. (A) Left panel shows Epithelial barrier function of *H. pylori*-infected polarized MKN28 cells as analyzed by measuring Trans-epithelial electrical resistance (TEER). Uninfected cells served as control. Y-axis represents % baseline resistance calculated as described in the experimental methods section. Right panel shows FITC-Dextran permeability values of *H. pylori*-infected MKN28 cells. Uninfected cells served as control. Y-axis represents the relative fluorescence units. *indicates p-value<0.05. (B) Confocal micrographs of 24 hr *H. pylori*-infected MKN28 cells showing delocalization of claudin-4 from the tight junctions as indicated by the white arrows (Lower panel). 3-dimensional (3D) reconstruction of images was performed using Imaris version 7.6. Uninfected cells served as control. Blue, DAPI-stained nuclei; Red, Cy3-stained claudin-4; Green, Alexaflour 488-stained *H. pylori*. (C) Western blots of membrane and cytoplasmic protein fractions of *H. pylori*-infected and uninfected MKN28 cells indicating localization of claudin-4. P-cadherin and beta-tubulin served as controls for membrane (Mem) and cytoplasmic (Cyto) fractions, respectively (upper panel). Western blots of cytoplasmic and nuclear protein fractions of *H. pylori*-infected and uninfected cells indicating localization of claudin-4 (lower panel). Beta-tubulin and Lamin A/C served as controls for cytoplasmic and nuclear fractions, respectively. The image is representative of 3 independent experiments. (D). Total protein lysate from *H. pylori*-infected and uninfected MKN28 cells at indicated time points were immunoblotted using claudin-4 antibody. β-actin serves as loading control for the experiment. The image is representative of 3 independent experiments (left panel). Total mRNA from *H. pylori*-infected and uninfected cells were analysed for claudin-4 gene expression using qPCR. The experiment was done in triplicate and the relative quantitation are expressed as histobars (right panel). GAPDH served as the endogenous control for the experiment. UN: uninfected; WT: *H. pylori-*infected.

In order to visualize the localization of claudin-4 in *H. pylori*-infected and uninfected cells, confocal laser scanning microscopy (CLSM) imaging was used. Within 24 hours of apical exposure of MKN28 cells to *H. pylori*, delocalization of claudin-4 from the cell-cell tight junctions was observed when compared to uninfected cells. Fig. 1B shows 3D reconstruction of the z-stack images of both *H. pylori*-infected and uninfected cells. The 3D images clearly display delocalization of claudin-4 from the tight junctions. *H. pylori*-infected cells showed cytoplasmic and peri-nuclear staining of claudin-4 whereas the uninfected cells showed localization of these proteins at the cell-cell junctions marking clear boundaries between cells (Fig. 1B).

To further support the finding that *H. pylori* affects the localization of claudin-4, western blot analysis was employed to examine the expression of claudin-4 in the membrane, cytoplasmic and nuclear protein fractions. The result shows an increased cytoplasmic and nuclear expression of claudin-4 suggesting delocalization of these proteins from cell-cell junction in *H. pylori*-infected cells. Furthermore, the membrane claudin-4 expression in *H. pylori*-infected cells was found to be reduced when compared to uninfected cells affirming the delocalization of claudin-4 (Fig. 1C).

We next examined the overall protein and mRNA expression of claudin-4 of uninfected cells and *H. pylori*-infected cells using western blot analysis and qPCR studies. We found that the protein expression did not differ between *H. pylori*-infected cells and uninfected cells at various time points studied (Fig. 1D, upper panel). Similarly, mRNA expression of claudin-4 in *H. pylori*-infected and uninfected cells did not differ (Fig. 1D, lower panel). The results indicate that *H. pylori* induced delocalization of claudin-4 is not dependant on the expression level.

### Inhibition of ERK activation minimized H. pylori-induced delocalization of claudin-4

It was reported that during *H. pylori* infection, increased levels of EGF-related peptides and NOD1 dependant mechanisms activate ERK^[34]^. Wang *et al* in 2004 demonstrated that ERK activation triggered barrier function disruption in human corneal epithelial cells^[35]^. We therefore proceeded to investigate whether inhibition of ERK activation using MEK inhibitor (U0126) would alter the process of *H. pylori*-induced barrier function disruption. Results show increased epithelial resistance (TEER) was detected in U0126-treated *H. pylori*-infected cells as compared to infected cells without inhibitor treatment (p-value = 0.003098682) (Fig. 2A, left panel). Interestingly, uninfected cells with or without inhibitor treatment showed similar TEER values. Furthermore, there was reduced permeability of 4kDa FITC-Dextran in *H. pylori*-infected cells with U0126 treatment as compared to infected cells without inhibitor treatment (Fig. 2A, right panel). The results show that upon the inhibition of *H. pylori*-induced ERK activation, the host epithelial barrier function is normalized. The data suggest that *H. pylori*-induced ERK activation has an important role in regulating epithelial barrier function in MKN28 cells supporting the findings of Wang et al (2004) demonstrated in corneal epithelial cells^[35]^.

**Fig.2.**
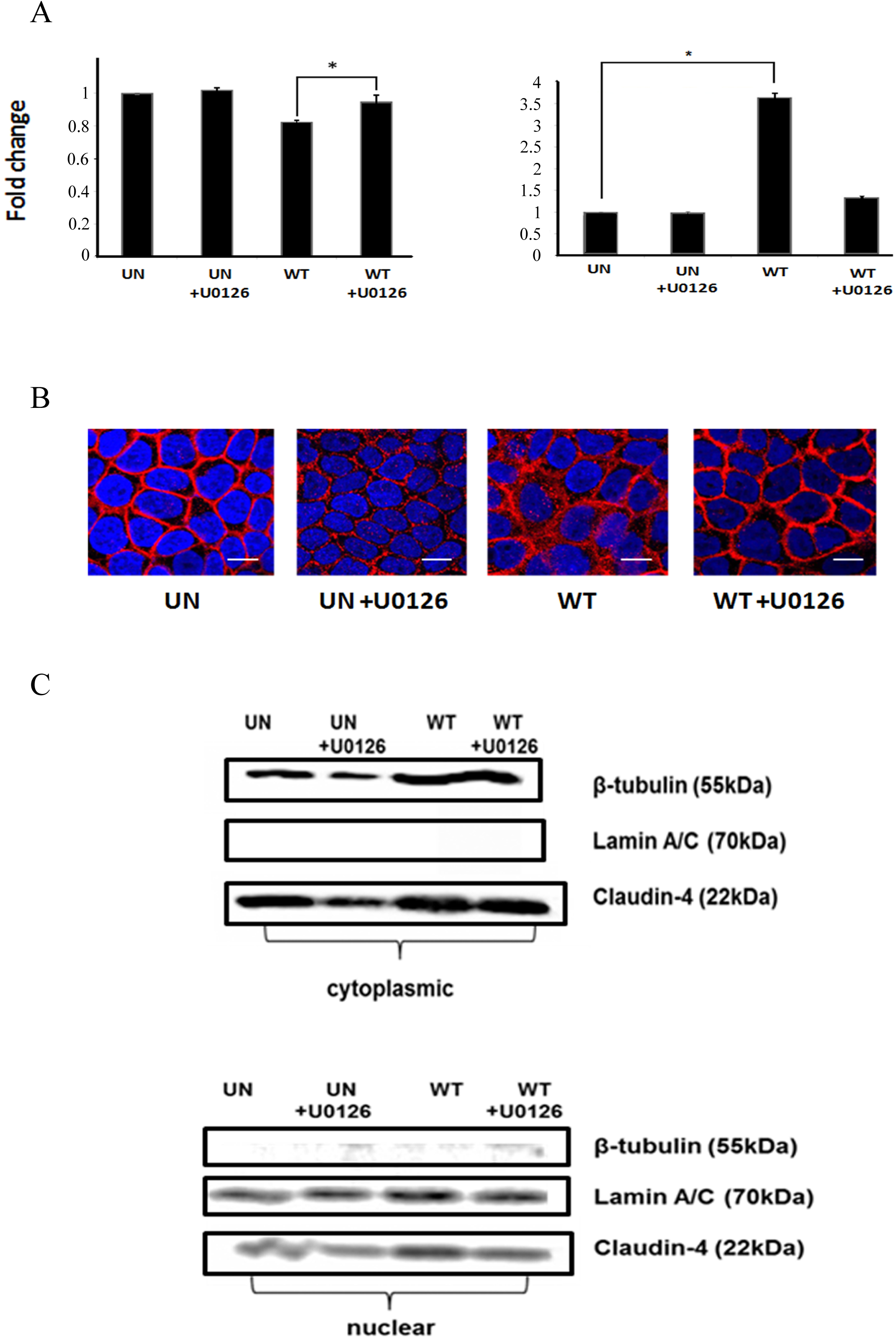
Effect of pretreatment with ERK activation inhibitor, U0126, on *H. pylori*-infected MKN28 cells. (A) Epithelial barrier function of *H. pylori*-infected cells with and without U0126 treatment was analyzed by measuring Trans-epithelial electrical resistance (TEER). Y-axis represents % baseline resistance calculated as described in the experimental methods section (left panel). FITC-Dextran permeability values of *H. pylori*-infected cells with and without U0126 (MEK-inhibitor) treatment were determined. *indicates p-value<0.05. Y-axis represents the relative fluorescence units (right panel). Uninfected cells with and without inhibitor served as controls. UN: uninfected, WT: wild-type *H. pylori*-infected. (B) Confocal micrographs of *H. pylori*-infected MKN28 cells with and without U0126 treatment showing localization of claudin-4. Blue, DAPI-stained nuclei; Red, Cy3-stained claudin-4. (C). Western blots of membrane and cytoplasmic protein fractions of *H. pylori*-infected cells with or without U0126 treatment indicating localization of claudin-4. (upper panel) P-cadherin and beta-tubulin served as internal controls for membrane and cytoplasmic fractions, respectively. Western blots of cytoplasmic and nuclear protein fractions of U0126 treated *H. pylori*-infected cells indicating the localization of claudin-4 (lower panel). Uninfected cells with and without inhibitor treatment served as controls. Beta-tubulin and Lamin A/C served as controls for cytoplasmic and nuclear fractions, respectively. Uninfected cells with and without inhibitor served as controls. UN: uninfected cells, WT: wild-type *H. pylori*-cells.

CLSM was used to further investigate the role of *H. pylori*–activated ERK in delocalizing claudin-4 in support of the TEER and FITC-Dextran results. Immuno staining of *H. pylori*-infected MKN28 cells treated with ERK activation inhibitor (U0126) showed minimized redistribution of claudin-4 when compared with *H. pylori*-infected cells without inhibitor treatment (Fig. 2B). More interestingly, western blots demonstrated that the redistribution of claudin-4 from membrane to cytoplasm and nucleus was clearly reduced in U0126-treated *H. pylori*-infected cells (Fig.2C).

### Transcriptomic changes induced by H. pylori infection leads to cell junction disruption

Our findings have thus demonstrated that *H. pylori* (infection?) plays an essential role in the disruption of epithelial barrier function with the involvement of ERK activation. In order to further elucidate the role of *H. pylori* in regulating signalling pathways and induce cell-cell junction disruption, RNA-Seq analysis was performed on *H. pylori*-infected primary gastric epithelial cells.

In this study, quality control analysis of the downloaded libraries was executed and the output showed that the mean quality score of the reads is ∼37 indicating high quality sequencing^[40]^. Following which, mapping of the raw reads was accomplished. The percentage of uniquely mapped reads was ∼92% with over 90% of those reads mapped to exons. This indicates the reliability of the libraries for further downstream analyses. Subsequently, differential gene expression analysis revealed that 8472 genes were significantly regulated in *H. pylori*-infected cells. Of these, 3665 genes were found to be upregulated while 4807 were downregulated (Fig. 3A). Further to this, the differential regulated genes as shown in Fig. 3B, were found to include interleukin-8 (IL8) and matrix metalloproteinase 10 (MMP10). These two genes have previously been shown to be upregulated in response to *H. pylori* infection^[46][47]^ further supporting the credibility of the differential gene expression analysis. The scatter plot also shows the upregulation of the genes that are associated with tight junction assembly and host signalling pathway (CLDN18, MAP2K1 and NFKB1A) (Fig. 3B).

**Fig.3.**
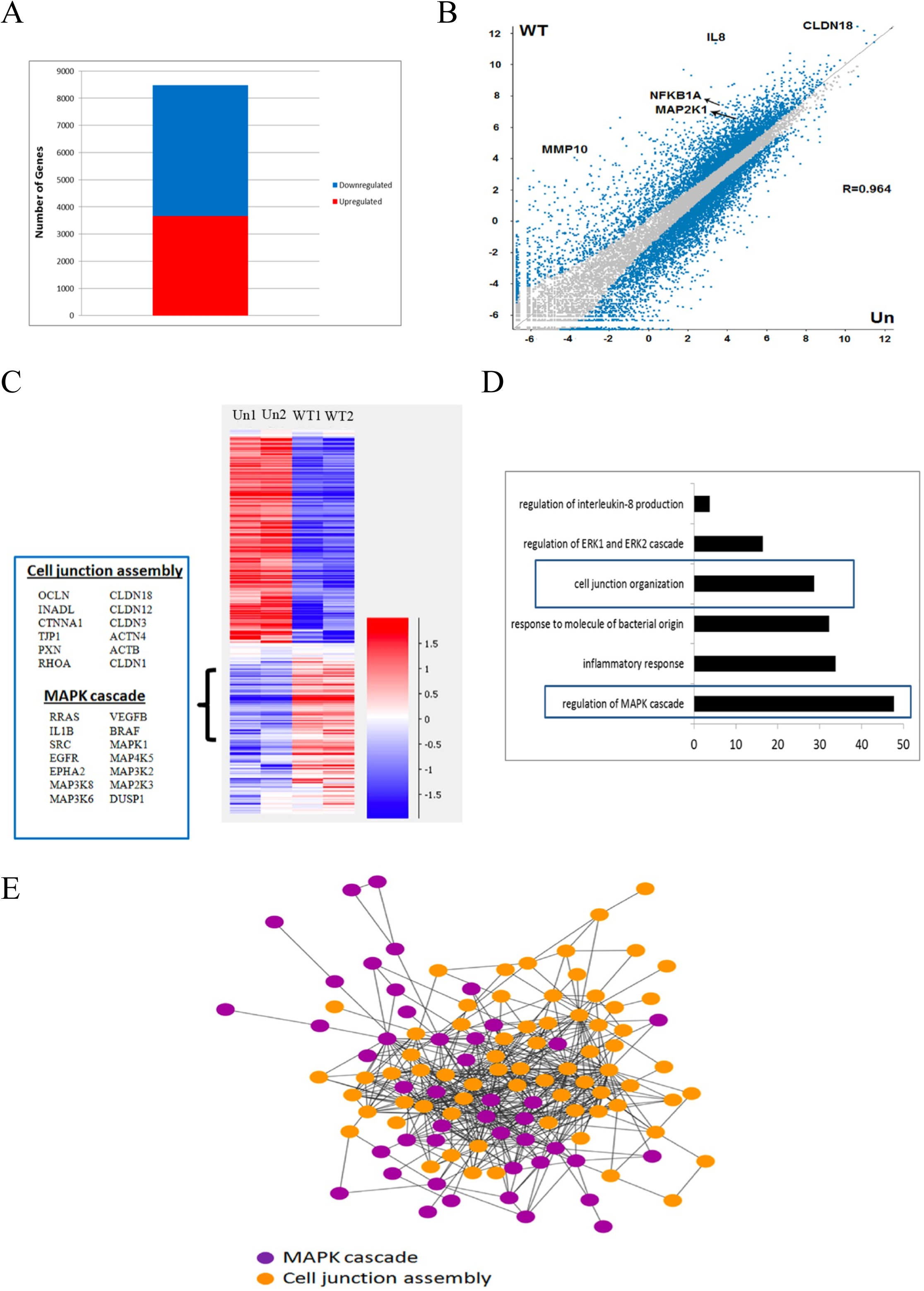
(A) Stacked column showing the number of significantly upregulated and downregulated genes in *H. pylori*-infected primary gastric cells. X-axis represents number of genes. (B) Scatter plot reveals the transcriptomic changes induced in *H. pylori*-infected cells. Grey dots represent genes that are not regulated by *H. pylori* whereas the blue dots represents the genes which are significantly altered in response to *H. pylori* infection. Representative dots of significantly upregulated genes are labelled on the plot. X-axis indicates the log10(RPKM) of the genes in uninfected cells (un) and Y-axis indicates the log10(RPKM) of the genes in *H. pylori*-infected cells (WT). (C) Heatmap demonstrating the RPKM values of the significantly regulated genes across replicates. The color scale depicts the log10(RPKM) starting from dark blue (lowest) to dark red (highest). The box to the left side presents examples of genes upregulated in *H. pylori*-infected cells and belonging to the indicated biological processes.UN1:Uninfected 1; UN2: Uninfected 2; WT1: *H. pylori*-infected 1; WT2: *H. pylori*-infected 2. (D) Gene Ontology analysis of genes upregulated in *H. pylori*-infected cells denotes the enrichment of the indicated biological processes terms. The cell junction and MAPK cascade associated GO terms are highlighted. X-axis is the – log10(p-value) of the enriched GO term. (E) Interactions among proteins that are upregulated in *H. pylori*-infected cells and associated with cell junction assembly and MAPK cascade are visualized using Cytoscape 3.2.0. The violet nodes depict the MAPK cascade associated genes and Orange nodes depict the cell junction assembly related genes.

The scaled Reads Per Kilobase Million (RPKM) values of all the significantly regulated genes were visualized across the replicates using a hierarchical clustered heatmap. Among the significantly upregulated genes (cutoff p-value = <0.05), we found many genes which are closely associated with tight junction assembly as well as MAPK cascade indicating that both the processes are heavily affected in *H. pylori*-infected cells. The heatmap also demonstrated similar expression profile among the replicates of each condition (Fig. 3C). Additionally, gene ontology analysis of *H. pylori* upregulated genes shows enrichment of cell junction assembly and MAPK cascade regulation processes. This has provided more evidence suggesting the role of *H. pylori* in the regulation of these two processes. Other processes such as response to bacterial proteins and inflammation were also observed to be enriched (Fig. 3D).

The upregulated genes associated with cell junction assembly and MAPK cascade were further subjected to protein-protein interaction analysis using STRING database and visualized using Cytoscape 3.2.0. Most astoundingly, the network reveals a strong protein-protein interaction among the upregulated genes involved in both the indicated biological processes forming an extremely tight network (Fig. 3.E). This result highly suggests the extent of gene regulation exerted by *H. pylori* on host cells where the bacterium induces disruption of cell-cell junction disruption via activation of MAPK/ERK associated pathways.

## Discussion

Cell-cell tight junctions, being at the apical surface, constitute an important protective barrier which when compromised can lead to a multitude of deleterious effects on the host. Thus, the proteins that are responsible for forming the tight junction assembly could be crucial targets for pathogens to initiate pathogenesis. Claudins, one of the two major tight junction protein complex, are highly expressed in gastric tissue and has been shown to have aberrant expression and localization in cancer conditions^[48]^. It has also been reported to be a multifunctional protein with many roles in cell migration, cell signalling and barrier maintenance^[49]^. In the event of infection and inflammation, internalization of claudins could lead to deleterious effects on the cell^[8]^. A recent study has shown that claudin-4 was over-expressed in 71% of 192 gastric cancer cases studied^[50]^. The importance of claudin-4 in normal function of gastric mucosal barrier should not be overlooked. Our study examines the course of claudins (in particular claudin-4) during *H. pylori* infection.

In this study, polarized MKN28 cells were used thanks to the organized cell–cell junctions^[29]^. The deleterious effect of *H. pylori* on disrupting the barrier function of MKN28 cells was shown by, the significant reduction in the TEER measurements and a concomitant increase in FITC-Dextran permeability in *H. pylori*-infected cells. Our findings strongly suggest that *H. pylori* induces delocalization of claudin-4 via activated ERK pathway leading to the loss of barrier function of the cell-cell tight junctions. The results support earlier reports that *H. pylori* impairs barrier function during pathogenesis^[29][51][52]^.

Our initial data suggest that claudin-4 is redistributed from the cell-cell tight junctions into the cytoplasm. This is in agreement with several earlier reports that suggested the delocalization of tight junction proteins as a marker for transformed cells^[9][11]^. But, Fedwick *et al*., (2005) reported that *H. pylori* SS1-infected non-transformed polarized SCBN cells showed both the disruption of cell-cell junction and the reduction of the total protein level of claudin-4 and -5^[28]^. In order to visualize the effect of *H. pylori* on host claudin-4, we used immunofluorescence imaging which clearly show the delocalization of claudin-4 from cell-cell tight junctions to the cytoplasm and nucleus in *H. pylori*-infected cells. Furthermore, there was no obvious display of diminished staining-density of claudin-4 (Fig. 1B). Our confocal finding is further supported by western blot analysis and qPCR data (Fig. 1D) that revealed the overall level of expression of claudin-4 did not alter. Furthermore, western blot analysis (Fig. 1C) showed an increase of claudin-4, an apically expressed protein, in the cytoplasmic fraction but a decrease in the membrane fraction in *H. pylori*-infected cells. Additionally, the expression of claudin-4 in the nuclear fraction was also found to increase in *H. pylori*-infected cells as compared to uninfected cells. Taken together, the findings in this study affirm that there is delocalization but not reduction in claudins in *H. pylori*-infected cells as reported earlier ^[28][53]^.

Activated ERK pathway has been reported to play a major role in inducing various host responses during pathogenesis^[33][34]^. Studies have reported that during *H. pylori* infection, increased levels of EGF and other related proteins have been shown to activate the EGFR pathway, signalling a cascade of events thereafter^[54]^. Of interest are recent reports that suggested activated ERK molecules have a role in epithelial barrier dysfunction^[35][55]^. By inhibiting ERK activation using U0126 in *H. pylori*-infected cells, there was significant reduction in the delocalization of claudin-4 (Fig.2B). Similarly, redistribution of claudin-4 into the cytoplasmic and nuclear region was reduced as compared to *H. pylori*-infected cells treated with U0126 (Fig.2C). Furthermore, the cell-cell barrier function in MKN28 cells was found to be maintained in cells treated with U0126 prior to infection (Fig.2A). It is therefore opportune to implicate that the activated ERK plays a role in delocalizing claudin-4 as a host response to *H. pylori* infection.

Our computational analysis demonstrated that *H. pylori* triggers the differential expression of many crucial genes including IL8 and MMP10 (Fig.3B) which is in agreement with the earlier reports that indicated these genes were regulated by *H. pylori*^[46][47]^. In addition to that, our RNA-Seq analysis revealed the upregulation of a large number of genes associated with MAPK cascade (Fig.3C). This is congruent with an earlier report that demonstrated the activation of MAPK cascade in *H. pylori*-infected cells, in a dose dependent manner^[56]^. However, our study is the first to report the extent by which *H. pylori* regulates the genes associated with the activation of MAPK cascade and cell junction assembly. Interactome analysis reveals that the upregulated MAPK cascade and cell junction genes interact closely with each other forming a tight and strong network (Fig. 3E). This asserts that the regulation of cell junction assembly by *H. pylori* could potentially be associated to the activation of MAPK cascade. This study reveals for the first time that *H. pylori-*activated ERK/MAPK cascade plays a significant role in disrupting cell-cell junctions in gastric epithelial cells.

This study has provided evidence suggesting *H. pylori*-activated ERK pathway plays a direct role in redistributing claudin-4 from the tight junctions resulting in compromising host barrier integrity. Interestingly, the transcriptomic analyses also revealed a wealth of information on the strong interactions at the molecular level between tight junction proteins and ERK signalling proteins. However, further analyses involving the genome binding occupancy profiling of transcription factors that are upregulated in *H. pylori*-infected cells are required to decipher the molecular mechanism by which *H. pylori* causes tight junction disruption via ERK activation.

